# Temporal instability of lake charr phenotypes: synchronicity of growth rates and morphology linked to environmental variables?

**DOI:** 10.1101/2020.08.13.249557

**Authors:** L. Chavarie, Steve Voelker, M.J. Hansen, C.R. Bronte, A.M. Muir, M.S. Zimmerman, C.C. Krueger

## Abstract

Pathways through which phenotypic variation arises among individuals arise can be complex. One assumption often made in relation to intraspecific diversity is that the stability or predictability of the environment will interact with expression of the underlying phenotypic variation. To address biological complexity below the species level, we investigated variability across years in morphology and annual growth increments between and within two sympatric lake charr ecotypes in Rush Lake, USA. We found a rapid phenotypic shift in body and head shape within a decade. The magnitude and direction of the observed phenotypic change was consistent in both ecotypes, which suggests similar pathways caused the temporal variation over time. Over the same time period, annual growth increments declined for both lake charr ecotypes and corresponded with a consistent phenotypic shift of each ecotype. Despite ecotype-specific annual growth changes in response to winter conditions, the observed annual growth shift for both ecotypes was linked, to some degree, with variation in the environment. Particularly, a declining trend in regional cloud cover was associated with an increase of early stage (age 1-3) annual growth for lake charr of Rush Lake. Underlying mechanisms causing reduced growth rates and constrained morphological modulation are not fully understood. An improved knowledge of the biology hidden within the expression of phenotypic variation promises to clarify our understanding of temporal morphological diversity and instability.

## Introduction

A rapidly changing climate can have wide-ranging effects on organisms across ecosystems, which fosters a need to understand how ecosystems will respond to this variation in terms of structure and function (Montoya José & Raffaelli 2010; Pacifici *et al*. 2015). Contemporary climate change, which includes rapid increases in global temperatures, represents one of the most serious and current challenges to ecosystems, not only by threatening ecosystems directly (Norberg *et al*. 2012), but also by contributing to cumulative and additive effects with other perturbations (e.g., industrial development, pollution, overhavest, non-native species; CAFF 2013; Poesch *et al*. 2016). Ecosystems are mosaics of different habitats, climate change combined with abiotic and biotic variation across these habitats may lead to major eco-evolutionary responses (Grimm *et al*. 2013; Ware *et al*. 2019). Rapid biological responses to variation associated with climate change have already been detected at all levels, from individuals to species, communities, and ecosystems (Heino *et al*. 2009).

The importance of phenotypic variability has been emphasised in evolutionary and ecological population dynamics (Kinnison & Hairston Jr 2007; Schoener 2011), because variation fuels evolutionary change (Stearns 1989). Pathways through which phenotypic variation arises among individuals can be complex. Intraspecific trait variation as expressed within phenotypes, such as morphology or growth, can affect population dynamics through reproductive and mortality pathways (Bolnick *et al*. 2011). Furthermore, the magnitude of plasticity in the variation of trait expression differs among populations and ecotypes within a population (Skúlason *et al*. 2019). A conceptual framework to predict evolutionary and ecological consequences of climate change is currently limited by the scarcity of empirical data demonstrating phenotypic changes over time among individuals within ecosystems. The causes, patterns, and consequences of ecological and evolutionary responses to differences induced by environmental variability need to be quantified across species, space, and time.

The effect of growth rate heterogeneity on morphology modulation (e.g., heterochonry, allometry; Klingenberg 2014) is not a mechanism fully understood, despite being observed to constrain or enhance morphological differences in several fish species (Olsson *et al*. 2006; Heino 2014; Jacobson *et al*. 2015). Variation in developmental rate associated with juvenile growth rates has been demonstrated to have an effect on the origin of some ecotypes (Alexander & Adams 2004; Helland *et al*. 2009; McPhee *et al*. 2015). Yet how variation in early development and juvenile growth rate influence later morphology remains ambiguous, with almost no attention focused on among-individual variation within an ecotype. Models have often assumed simple population (and ecotype) dynamics, with populations (and ecotypes) commonly showing a stable ecological evolutionary equilibrium (Svanbäck *et al*. 2009; Skúlason *et al*. 2019).

Persistence of phenotypes within a system is generally associated with a relatively stable ecological environment that favors canalized phenotypes (e.g., reduction of phenotypic variance at either the within-genotype-among-individual or within-individual levels) over plastic phenotypes (Schluter 2000; Nosil 2012; Westneat *et al*. 2015; Skúlason *et al*. 2019). The propensity for growth has both environmental and heritable components with variability in phenotypic expression (van der Have & de Jong 1996; Kingsolver *et al*. 2004), even for canalized phenotypes. Given rapid environmental change occurring within aquatic ecosystems, phenotypic changes among individuals must not be taken as negligible, because phenotypic variation is driven by switches along developmental pathways (West-Eberhard 2003), in cases where developmental system flexibility can adjust immediately to environmentally variable conditions (Japyassú & Malange 2014). Thus, temporal variation in phenotypes within an ecotype may not be ecologically trivial in a rapidly changing world.

Records of organismal growth patterns provide long-term data sets that reflect environmental variation (Pereira 1995). The diversity of species and ecosystems for which growth chronologies exist (e.g., trees, bivalves, corals, fishes; Black 2009) provide robust datasets for assessment of temporal and spatial environmental variation and its ecological consequences across ecotypes and species, trophic levels, and communities that link ecosystems and biomes. The lake charr (*Salvelinus namaycush*) is a suitable fish species to link growth chronologies with environmental variation because their longevity that can exceed 60 years (Smith *et al*. 2008; Chavarie *et al*. 2016). Lake charr are also known to display intraspecific variation, mainly diversifying along a depth gradient, with shallow-versus deep-water ecotypes exploiting different prey resources within a lacustrine system (Chavarie et al. accepted). By selecting a case of co-existing shallow- and deep-water ecotypes of lake charr in Rush Lake (Fig. A1; Chavarie *et al*. 2016), we examined the response of lake charr to environmental variation below the species level. The effect of environmental variation, such as for temperature, on growth rates of aquatic organisms varies, especially across depths, in part because deep-water habitats are usually more environmentally stable (Thresher *et al*. 2007; Murdoch & Power 2013; Jeppesen *et al*. 2014). By exploiting different prey items (Chavarie *et al*. 2016), invertebrates (deep-water ecotype) vs. fish (shallow-water ecotype), lake charr ecoytpes in Rush Lake are likely to integrate diverse signals of climate variation across multiple trophic levels (Black *et al*. 2013a; Black *et al*. 2013b). Finally, Rush Lake is located near the southern edge of the lake chair’s species range in North America where variation in growth rate would likely be more sensitive to climate variation than for more populations closer to the center of the range and thereby may yield useful predictive responses to climate change.

In this study, we tested whether phenotypic expression of two Rush Lake lake charr ecotypes remained stable or changed over time. We predicted that if a rapid change in phenotypic expression occurred and this change was associated with environmental variation, the shallow-water ecotype would display a greater magnitude of variation than the deep-water ecotype due to the shallow-water habitat being more responsive and sensitive to contemporary climate change. We measured phenotypic expression in terms of morphology and growth chronology. The objectives of our study were to: 1) quantify the morphological variation between and within lake charr ecotypes over a ten-year period, 2) determine if patterns of growth chronologies expressed by ecotypes were temporally synchronous with each other and associated with the phenotypic variation displayed within each ecotype; and 3) examine if annual growth rates of lake charr ecotypes were related to environmental variation by using tree-ring cross-dating techniques.

## Material and Methods

Rush Lake is a small lake (1.31 km^2^) that contains deep-water (>80m) and is less than 2 km from Lake Superior (Chavarie *et al*. 2016). Rush Lake provided the first documented example of sympatric lake charr ecotypes in a small lake, diverging along a depth gradient. Two co-existing ecotypes of lake charr were found; a large streamlined-bodied shallow-water lake charr (lean ecotype) and a small plump-bodied deep-water ecotype (huronicus ecotype) analogous to the humper lake charr ecotype from Lake Superior (Hubbs 1929; Chavarie *et al*. 2016). The lean ecotype has a long head, long maxillae, and a posterior eye position, which are all characteristics for piscivorous feeding (Proulx & Magnan 2004; Keeley *et al*. 2007; Janhunen *et al*. 2009). The huronicus ecotype, with smaller gape and higher eye position than the lean ecotype, appears adapted for low-light vison and as a vertical migrating predator feeding on the invertebrate opossum shrimp (*Mysis spp.*) as its main prey (Hrabik *et al*. 2006; Muir *et al*. 2014).

### Assignment of lake charr ecotypes

Lake charr ecotypes were caught with bottom-set gillnets set in June 2007 and September 2018. Gillnets were deployed at depths from 10 to 83 m. Sets were made using 183 m long by 1.8 m high multifilament nylon gillnets consisting of stretch-mesh sizes from 50.8 to 114.3 mm, in 12.7 mm increments. Date, time, GPS location, and minimum and maximum water depth were recorded for each net set. All fish caught were photographed in lateral view (Muir *et al*. 2012) and sagittal otoliths were removed for analysis of age and growth.

Lake charr from each year were morphologically assessed and assigned to ecotype according to methods described by Muir *et al*. (2014). Lake charr caught in 2007 were previously assigned to ecotype (Chavarie *et al*. 2016) and assignments for lake charr caught in 2018 used the same methodology. Twenty sliding semi-landmarks and six homologous landmarks were digitized from each image to characterize head shape and 16 homologous and four sliding semi-landmarks were digitized from whole-body images to characterize body shape. Landmarks and semi-landmarks were digitized as x and y coordinates using TPSDig2 software (http://life.bio.sunysb.edu/ecotype). Digitized landmarks and semi-landmarks were processed in a series of Integrated Morphometrics Programs (IMP) version 8 (http://www2.canisius.edu/;sheets/ecotypesoft), using partial warp scores, which are thin-plate spline coefficients. Morphological methods and programs are described by Zelditch *et al*. (2012) and specific procedures were described by Chavarie *et al*. (2013). All morphological measurements were size-free by using centroid sizes (Zelditch *et al*. 2012). Ecotypes were assigned to each individual using a combination of Bayesian cluster analyses using head and body shape information (MCLUST; Fraley & Raftery 2009) and a visual identification by experienced lake charr biologists (M.J. Hansen & C. C. Krueger). Disagreements between model and visual assignments were settled using decision rules described by Muir *et al*. (2014).

### Temporal morphological variation between ecotypes and years

All data from 2007 and 2018 were combined to align all samples in the same shape space and partial warps for temporal morphological analyses between ecotypes and years. Principal component analysis (PCA) of body- and head-shape data were used to visualize morphological variation between and within lake charr ecotypes and years using PCAGen8 (IMP software). Canonical variate analyses (CVA) and validation procedures on body and head shape data were used to assess temporal differences within and between ecotypes (ecotype*year) using CVAGen (IMP software). Jacknife validation procedures included a test of assignment, with 1000 jackknife sets using 20% of our data as unknowns (Zelditch *et al*. 2012). Single-factor permutation multivariate analysis of variance (MANOVA) with 10000 permutations of CVAGen was used to test whether body and head shape differed between and within (i.e, years) ecotypes. If MANOVA indicated differences among groups (α < 0.05), procrustes distance means were calculated for pairwise comparisons using TWOGROUP from the IMP software as post-hoc tests (García-Rodríguez *et al*. 2011). Procrustes distance for each pairwise comparison described the magnitude of difference between ecotypes and years. A bootstrapped Goodall’s F statistic (*N* = 4900 bootstraps; full Procrustes based) was used to determine if pairwise comparison differed. Allometric trajectories in body and head shape were compared between ecotypes and years by regressing PC1 scores (size-free data) against centroid size (i.e., measure of size) (e.g., variation in developmental pathways can result in allometric trajectory patterns that can be parallel, divergent, convergent, or common; Simonsen *et al*. 2017); an allometric relationship occurred if the slope differed from 0.

Relative body condition was compared between ecotypes and years in a 2-way analysis of variance (ANOVA), with main effects for ecotype and year and the interaction between ecotypes and years (Zar 1999). To correct for size-related trends in body condition, relative body condition was defined as residuals from the power relationship between log10 (W) and log10 (TL). If the ecotype*year interaction was significant, years were compared within ecotypes and ecotypes were compared within years in 1-way ANOVAs. To visualize the results, least-squares means (+ SE) from the ANOVA were back-transformed from logarithms into original units of measure.

### Age assignment and growth increments from otoliths

Sagittal otoliths were used to estimate lake charr age and growth increments for fish sampled in 2007 and 2018. Otolith thin sections have been validated for age estimation of lake charr to an age of at least 50 years (Campana *et al*. 2008). Lake charr otoliths were prepared as described by Campana *et al*. (2008). One otolith from each fish was embedded in epoxy. A Buehler Isomet 1000 Precision Saw was used to remove a thin transverse section (400 μm) containing the nucleus perpendicular to the sulcus. Sections were mounted on glass slides and polished. Digital images of otolith sections were captured for age and growth assessment. Criteria for annulus demarcation followed those of Casselman and Gunn (1992). Age estimates were used to inform demarcation of growth increments, measured from the nucleus to the maximum ventral radius of the otolith, and radial measurements at each annulus were used to back-calculate length-at-age using the biological intercept back-calculation model (Campana 1990). The biological intercept (sagittal otolith radius = 0.137 mm; age-0 lake trout length = 21.7 mm; Hansen *et al*. 2012) was based on equations describing relationships between length, age in days, and sagittal otolith width of age-0 lake charr (Bronte *et al*. 1995).

### Back-calculated length at age from otoliths: growth patterns displayed by ecotypes through time

Otolith growth measurements can be used for several different purposes to gain ecological insight, but often need different analytical techniques to answer different questions. Towards this end, we used three analytical techniques and have provided a summary of the advantages and disadvantages of each (Table A1). To determine if patterns of growth chronologies expressed by ecotypes were temporally synchronous with each other, growth in length with age was modeled using a parameterization of the Von Bertalanffy length-age model (Gallucci & Quinn 1979; Quinn & Deriso 1999):

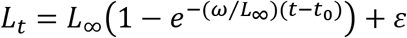

The length-age model describes back-calculated length *L_t_* (mm) at age *t* (years) as a function of age at length = 0 (*t*_0_ = years; incubation time of embryos from fertilization to hatching), early annual growth rate (ω = L_∞_×K = mm/year; Gallucci & Quinn 1979), theoretical maximum length (*L*_∞_ = mm), and residual error (*ε*). Parameters *ω* and *L*_∞_ were estimated using nonlinear mixed-effect models, package ‘nlme’ (R Core Team 2016), with a fixed population effect (the average growth curve for the population from which individual fish were sampled), individual as random effect (growth curves for individual fish sampled from the population), and sex (male or female), ecotype (lean or huronicus), or sampled year (2007 or 2018) as fixed factors (to compare average growth curves between sexes, ecotypes, and years; Vigliola & Meekan 2009). Mixed-effects models are appropriate for modelling the within-group correlation of longitudinal, auto-correlated, and unbalanced data, such as back-calculated growth histories (Pinheiro & Bates 2000). Five models of varying complexity were compared using AIC statistics (Burnham and Anderson 2002): (1) ecotypes and sample years both included, (2) ecotypes only included, (3) years only included, (4) sexes only included, and (5) neither ecotypes nor years included.

Annual growth increments were modeled using a linear mixed-effects model (Weisberg *et al*. 2010), wherein annual growth increments were modeled as a function of a fixed age effect (age of the fish when the increment formed), a random year effect (year in which the increment formed), a random fish effect (unique identifier for individual fish), and residual variation. The fixed age effect accounts for the fact that growth increments decline with age approximating a negative exponential curve. The random year effect reflects average increment width associated with each year of growth, after accounting for age effects (i.e., growth increment declines with age). The random year effect accounts for year-to-year environmental effects as random draws from a normal distribution, with a different draw for each year. The random fish effect accounts for fish-to-fish variation in growth as random draws from a normally distributed population with a different draw for each fish. The last source of variation is unmodeled residual variation. Differences between sexes (male or female), ecotypes (lean or huronicus) and sample years (2007 or 2018) were tested by including sex, ecotype, and sample year as fixed effects.

### Otolith increment cross-dating: growth of lake charr in relation to environmental variation

In recent decades, advancements in dendrochronological techniques have been increasingly applied to sagittal otoliths, leading to novel insights on how broad-scale climate variation can impact both freshwater and marine systems (Black *et al*. 2005; Matta *et al*. 2010; Black *et al*. 2013b). The use of dendrochronological methods (i.e., cross-dating techniques) ensures that specific growth annuli are assigned to an exact year (Black *et al*. 2005; Black *et al*. 2012; Black *et al*. 2013a). In turn, this process enhances connection to common environmental signals across fish, which have syncronously limited growth in certain years. To assign otolith growth increments to exact calendar years, transverse sections of sagittal otoliths were aligned by calendar year and cross-dated visually using the list method (Yamaguchi 1991) to confirm years when particularly large or small otolith increments would be expected. Thereafter, visual cross-dating was statistically confirmed using COFECHA software (Holmes 1983). In using COFECHA, otolith time-series with series intercorrelation values (i.e., Rbar) lower than 0.20 with the initial master chronology were removed and placed in a separate group that included more than one third of all fish sampled. In a previous study of lake charr otolith variation, Rbar values ranged from 0.42 to 0.97 (Black *et al*. 2013a), supporting the assumption that otolith width series with Rbar < 0.20 included anomalous inter-annual growth variation that did not match the initial master chronology. Otolith increment time series with Rbar < 0.20 underwent a second round of cross-dating with COFECHA, separately from those that matched better with the initial master chronology. Because otolith increment data grouped together by ecotype without *a priori* knowledge of ecotype, all subsequent chronologies and analyses were conducted separately based on ecotype assignment from the morphological analyses described above.

Dendrochronological detrending methods generally attempt to remove growth variation and emphasize inter-annual variation in growth controlled by climate (Fritts 1976). The first approach to detrending used the ARSTAN program (Cook and Krusic 2014) to fit cubic splines of various rigidity based on fish age. Thereafter, autoregressive modeling was used to enhance inter-annual growth variability and the resulting indexed time-series averaged within calendar years using a bi-weight robust mean. Thereafter, we undertook a second regional chronology standardization (RCS) approach known to better enhance low frequency signals compared to the cubic spline method (Table A1; Briffa & Melvin 2011). This detrending method divided each raw growth increment value by that expected from the mean growth increment for each ecotype and age combined (Fig. A2). These ratios were then multiplied by 100 and averaged within calendar years to yield a percentage change in growth for each year. In subsequent analyses of environmental influence on growth chronologies, data from each lake charr were truncated to feature only growth during young ages (age 1-3); which were excluded in earlier age-effect analyses but are a critical stage to phenotypic variation linked to environmental differences (Angilletta *et al*. 2004; Georga & Koumoundouros 2010; Ramler *et al*. 2014). The approach employed for these comparisons corrected for age directly rather than using ARSTAN detrending (see Methods above and Table A1). We also limited the analysis to calendar years where each combination of ecotype and collection period included otolith data from at least seven fish. This constrained the calendar years investigated to 1986 to 2012 for huronicus and 1988 to 2010 for leans.

Otolith increments, detrended with the ARSTAN program, were calculated as means within a year for both ecotypes and were initially compared against monthly resolution climate data for the corresponding year. Based on *a priori* knowledge of fish biology and lake-effect climate phenomena, temperature, precipitation, and cloud cover were selected as environmental variables (Chavarie *et al*. 2018; Voelker *et al*. 2019). Interpolated climate data (e.g., air temperature and precipitation) were obtained from ClimateNA version 5.6 software (Wang *et al*. 2016). Cloud cover climate data from airports within 7 km of Lake Superior were obtained at daily resolution from the NOAA Great Lakes Environmental Research Laboratory (https://www.glerl.noaa.gov/) and summarized by month.

Pearson correlation values were calculated between each ecotype-specific growth chronology and monthly climate variables for the corresponding and previous two year (i.e., to detect lag effect). After initial inspection of correlations between annual growth increments and monthly climate variables, the number of potential explanatory variables were consolidated into seasonal means, whereas winter to spring was defined as December through the next April for a corresponding-year, summer was defined as June to August, and fall was defined as September to November. Autocorrelation was expected to be present in otolith increment time series and resulting chronologies due to year-to-year lags in growth owing to fat storage and subsequent metabolic withdrawals, skip spawning effects, and climate and climatic-effects on water temperature. Robust assessments and modeling of autocorrelation on short time series is statistically impossible. Thus, we quantified what proportional weighting of climate data among the corresponding and two previous years produced the largest gains in Pearson correlations between otolith growth increment and seasonal climate data from an individual year to provide a window into how climate signals are incorporated into fish growth.

The influence of weighted seasonal variables on otolith growth increment was assessed using forward selection multiple regression models, package “lm” (R Core Team 2016). Final regression models were constructed for each ecotype separately to determine which individual seasonal climate variables as well as combinations of two or more seasonal variables were significant (α < 0.05) and resulted in higher Akaike Information Criterion values (Burnham & Anderson 2004).

## Results

### Temporal morphological variation between ecotypes and years

In total, 107 lake charr were sampled, including 39 huronicus and 20 leans in 2007 and 27 huronicus and 21 leans in 2018. For both ecotypes, lake charr caught in 2007 had deeper bodies than lake charr caught in 2018 (Fig. 1). The first two principal components explained 48.9% of the variation in lake charr body shape from Rush Lake (Fig. 1a). Body shape differed between years within each lake charr ecotype (CVA, Axis 1 ƛ = 0.015, *p* < 0.01 and Axis 2 ƛ = 0.29, *p* < 0.01; Fig. 1b). Jackknife classification of body shape had a 54.3% rate of correct year and ecotype assignment (i.e., ecotypes and years as different factors). Body shape means differed between ecotypes and years (Permutation MANOVA, F = 11.7, df = 3, *p* ≤0.01), and the magnitude of these differences was slightly larger between ecotypes than between years. Pairwise body shape comparisons differed between ecotypes for both 2007 and 2018 (F-tests; *p* ≤ 0.05). For lean and huronicus lake charr sampled in 2007, the Goodall’s F was 21.1 and distance between means was 0.030 ± 0.0028 (SE), whereas for the lean and huronicus sampled in 2018, the Goodall’s F was 18.2 and distance between means was 0.032 ± 0.0031 (SE). Pairwise body shape comparisons also differed between years for both ecotypes (F-tests; *p* ≤ 0.05). For huronicus 2007 vs 2018, the Goodall’s F was 10.8 and distance between means was 0.020 ± 0.0012 (SE), whereas for the lean 2007 vs 2018, the Goodall’s F was 10.4 and distance between means was 0.025 ± 0.0024 (SE). Allometric trajectories in body shape did not differ between 2007 and 2018, except for leans sampled in 2018 (R^2^ = 0.56, p ≤ 0.01).

**Fig. 1.**
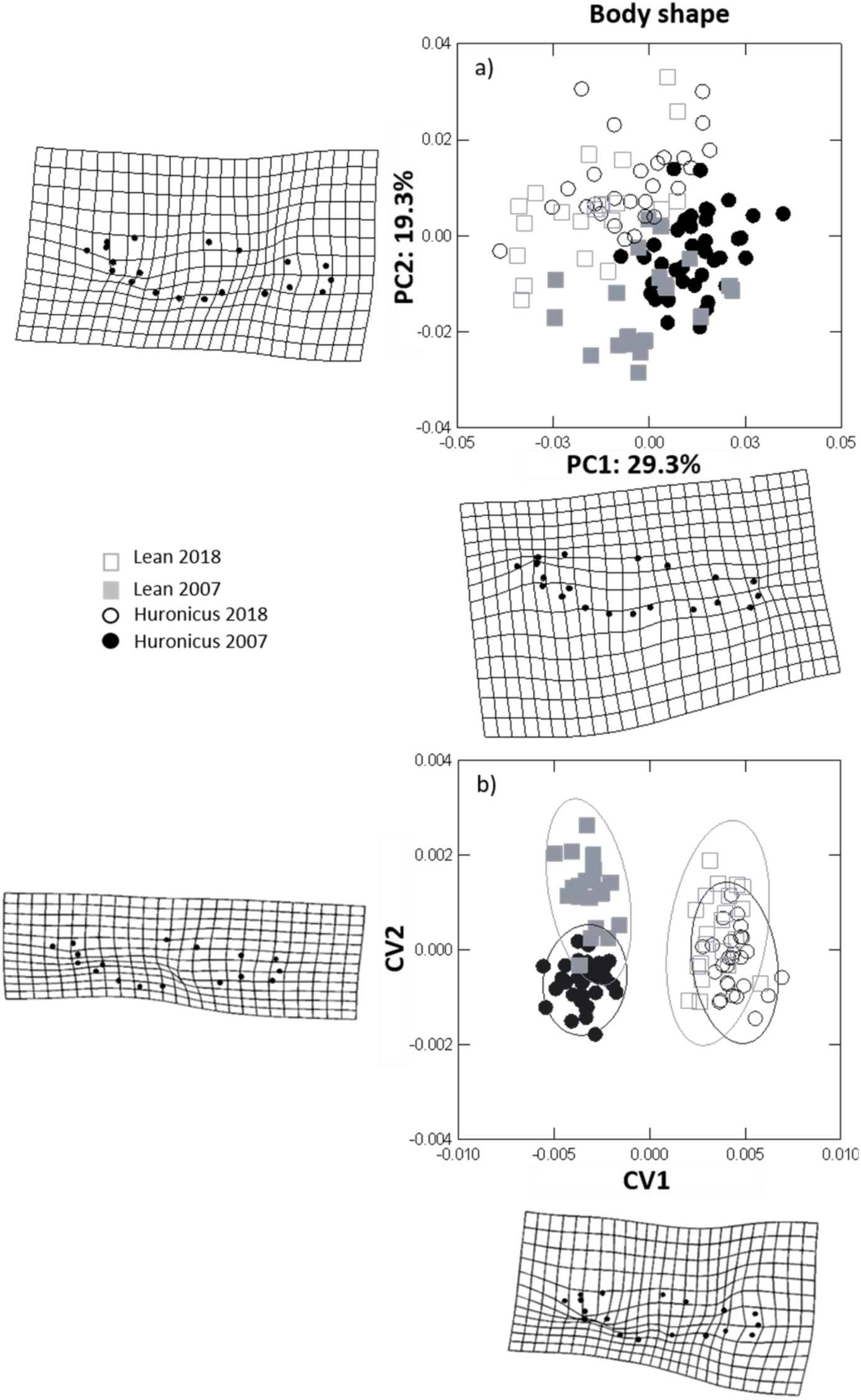
a) PCA of lake charr body shape with percentage representing the variation explained by that component and b) CVA of lake charr body shape with 95% confidence ellipses delineating groups. Deformation grid plots represent the body and head shape variation of lake charr sampled in Rush Lake in 2007 and 2018, in a positive direction, along each axis.

For both ecotypes, lake charr from 2007 had longer and deeper heads than lake charr from 2018 (Fig. 2). The first two principal components explained 61.2% of the variation in lake charr head shape (Fig. 2a). Head shape differed between years within each lake charr ecotype (CVA, Axis 1 ƛ = 0.012, *p* < 0.01 and Axis 2 ƛ = 0.13, *p* < 0.01; Fig. 2b). Jackknife classification on head shape had a 49.5% rate of correct year and ecotype assignment (i.e., ecotypes and years as factors). Head shape differed between ecotypes and years (Permutation MANOVA, F = 21.75 df = 3, *p* ≤ 0.01; Fig 3), and the magnitude of these differences was slightly larger between years than between ecotypes. Pairwise comparisons of head shape differed between ecotypes for both 2007 and 2018 (F-tests; *p ≤* 0.05). For lean and huronicus sampled in 2007, Goodall’s F was 21.7 and distance between means was 0.040 ± 0.0035 (SE), whereas for the lean and huronicus sampled 2018, Goodall’s F was 21.0 and distance between means was 0.048 ± 0.0044 (SE). Pairwise head shape comparison also differed between years for both ecotypes (F-tests; *p ≤* 0.05). For the huronicus, Goodall’s F was 54.9 and distance between means was 0.063 ± 0.0036 (SE), whereas for the lean, Goodall’s F was 38.6 and distance between means was 0.063 ± 0.0041 (SE). Allometric trajectories in head shape did not differ except for lean lake charr in 2018 (R^2^ = 0.36, *p* < 0.01; Fig. 3).

**Fig. 2.**
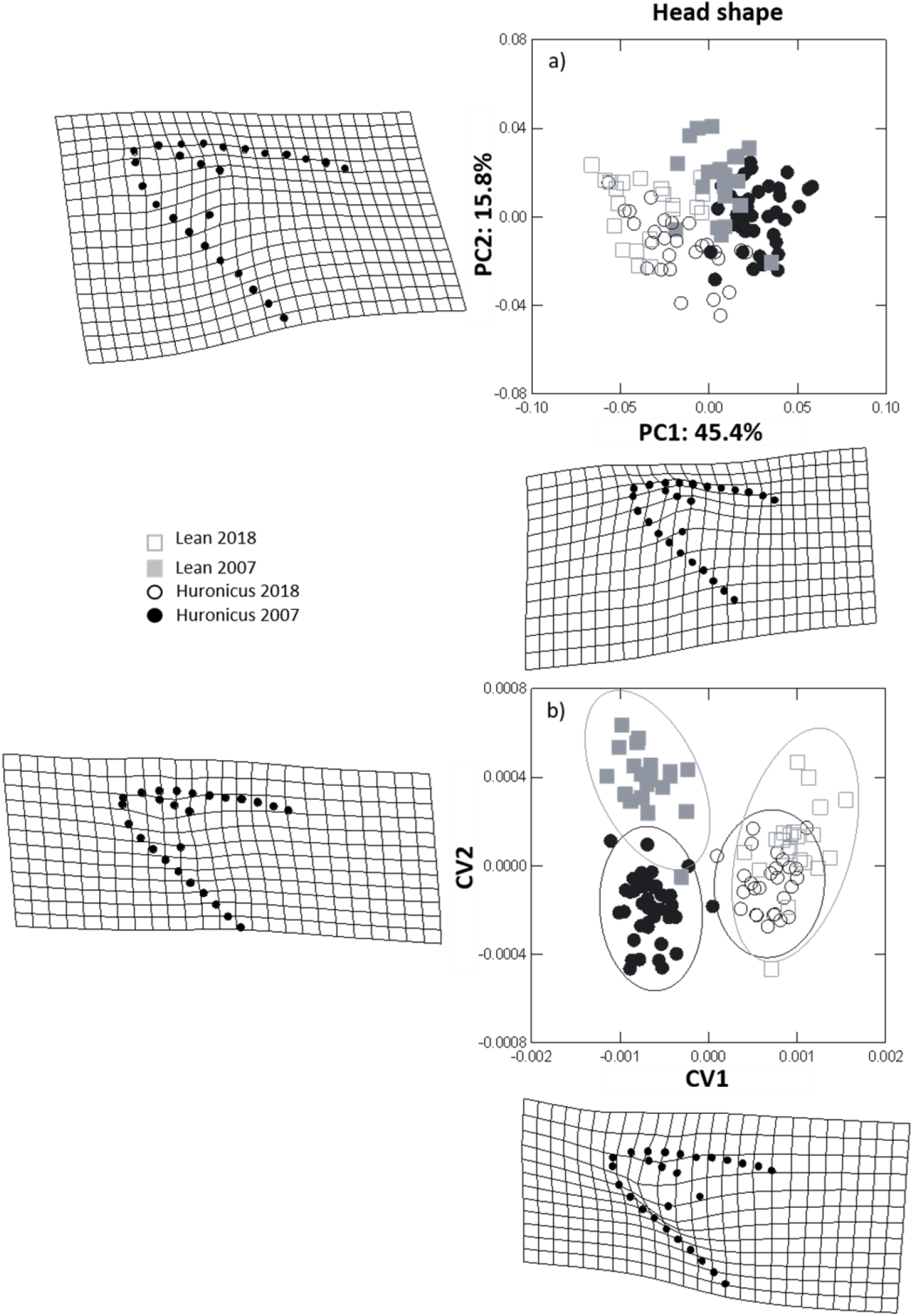
a) PCA of lake charr head shape with percentage representing the variation explained by that component and b) CVA of lake charr head shape with 95% confidence ellipses delineating groups. Deformation grid plots represent the body and head shape variation of lake charr sampled in Rush Lake in 2007 and 2018, in a positive direction, along each axis

**Fig. 3.**
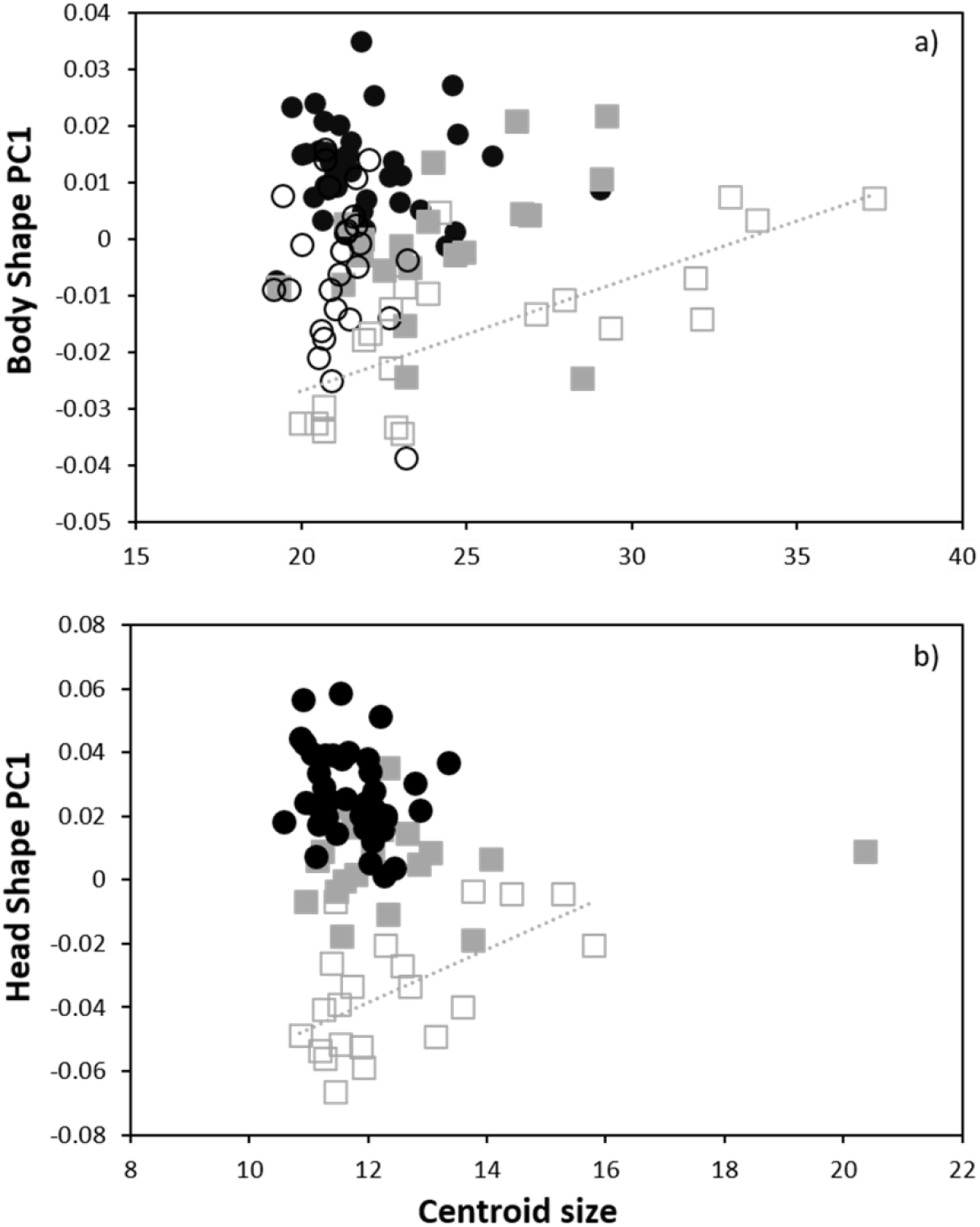
Body and head shape changes through growth of lean and huronicus lake charr ecotypes from 2007 and 2018. PC1 scores of body shape (a) and head shape (b) are plotted against centroid size. Only lean 2018 regressions were significantly different from 0, for both body and head shape (body shape: R^2^ = 0.56, p≤0.01; head shape: R^2^= 0.36, p≤0.01). Lean are represented by squares and huronicus by circles, whereas filled symbols are individual sampled in 2007 and empty symbols are lake charr caught in 2018.

Relative body condition differed between ecotypes and years (ecotype*year: *F*_1, 134_ = 15.0; *P* < 0.01; Fig. 4.). Within years, relative body condition of the huronicus ecotype was higher than the lean ecotype in 2007 (*F*_1, 66_ = 25.4; *P* < 0.01) but similar to the lean ecotype in 2018 (*F*_1, 68_ = 0.2; *P* = 0.7). Within ecotypes, relative body condition of the huronicus was higher in 2007 than in 2018 (*F*_1, 80_ = 55.1; *P* < 0.01) but the lean ecotype did not differ between years (*F*_1, 54_ = 0.4; *P* = 0.5).

**Fig. 4.**
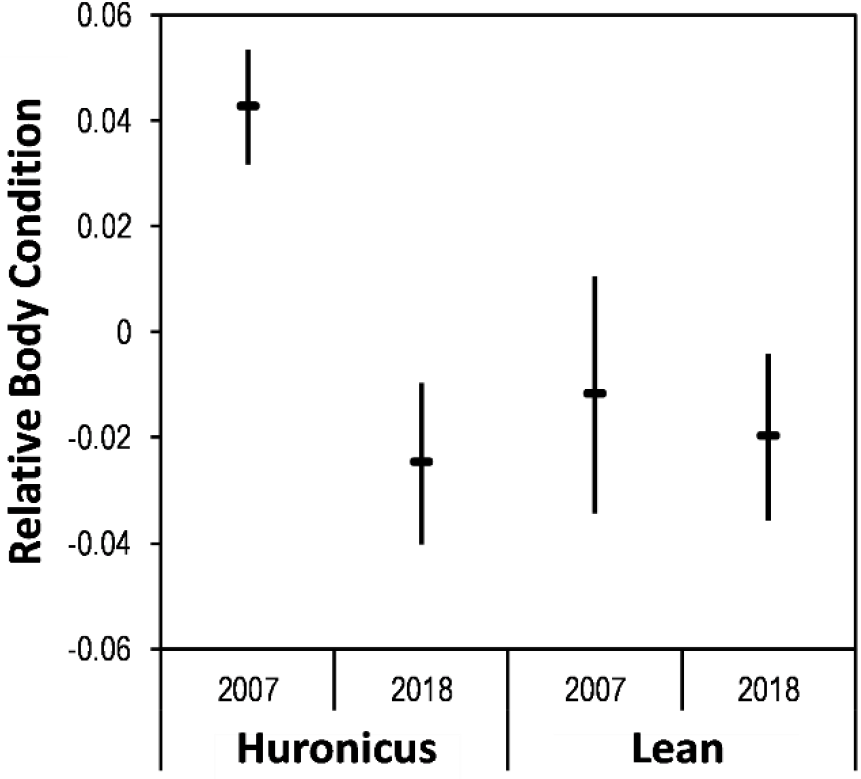
Relative body condition of huronicus and lean lake charr sampled from Rush Lake, in 2007 and 2018.

### Otoliths back-calculated growth: growth patterns displayed by ecotypes through time

Length-at-age of lake charr was best described by separate models for ecotypes and sample years (Table 1). Lean lake charr grew faster to a longer asymptotic length than huronicus lake charr in 2007 and 2018 (Fig. 5). Lean and huronicus lake charr sampled in 2018 grew faster at early age than those sampled in 2007, whereas both ecotypes sampled in 2007 grew to longer asymptotic length than those sampled in 2018. The early growth rate of lean lake charr sampled in 2007 was similar rate to huronicus lake charr sampled in 2018. In contrast, the asymptotic length of lean lake charr sampled in 2018 was similar to the asymptotic length of huronicus lake charr sampled in 2007.

**Table 1.** Tests of fixed effects for differences between lake charr ecotypes (lean, huronicus), sample years (2007, 2018), and sexes (male, female) from a nonlinear mixed-effects model of back-calculated length at sagittal otolith age, with random fish effects (fish-to-fish variation in growth) sampled in Rush Lake.

| Models | df | logLik | AIC | $\Delta_i$ | $e^{(-0.5*\Delta_i)}$ | $w_i$ |
| --- | --- | --- | --- | --- | --- | --- |
| Ecotypes + Years | 19 | -7672.0 | 15382.0 | 0.0 | 1 | 1 |
| Ecotypes | 13 | -7702.4 | 15430.8 | 48.8 | 0 | 0 |
| Years | 13 | -7705.7 | 15437.4 | 55.4 | 0 | 0 |
| Sexes | 13 | -7729.0 | 15484.0 | 102.0 | 0 | 0 |
| Null | 10 | -7732.9 | 15485.8 | 103.8 | 0 | 0 |

**Fig. 5.**
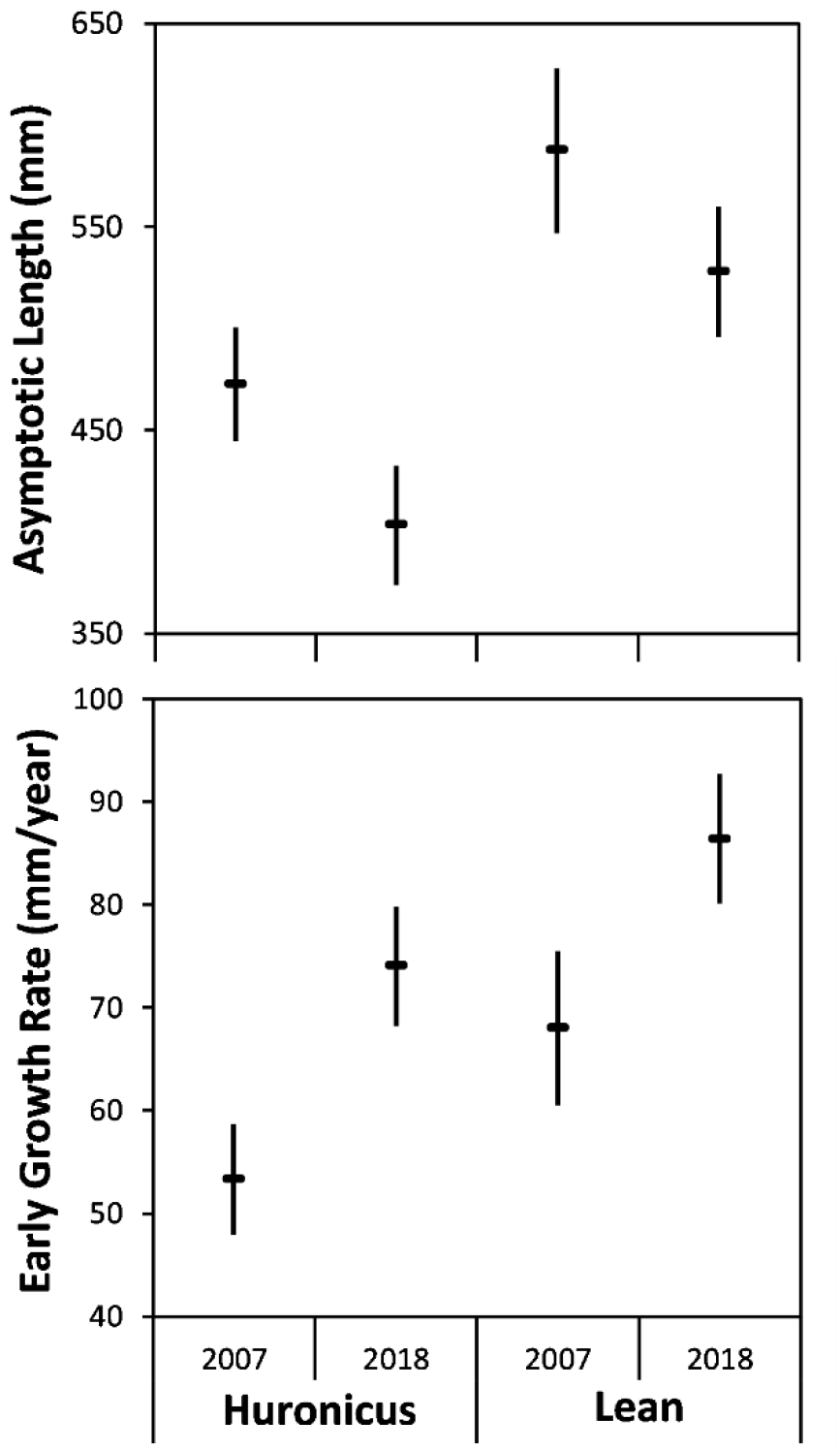
Asymptotic length (mm) and early growth rate (mm/year; first year) calculated from sections sagittal otoliths of huronicus and lean lake charr sampled from Rush Lake, in 2007 and 2018.

Average otolith growth increments (corrected for age) differed between lean and huronicus ecotypes (*F*_1, 1927_ = 16.9; *P* < 0.01), but not between males and females (*F*_1, 1927_ = 0.4; *P* = 0.52; Table 2) or between sampling years (*F*_1, 1927_ = 0.08; *P* = 0.77; Table 2). Average annual growth increments of huronicus and lean lake charr fluctuated without a specific trend prior to calendar year 2009 and then declined steadily between 2009 and 2018 (Fig. 6). For huronicus lake charr, average annual growth increments varied without temporal trend from 1977 until 1988, increased slowly and erratically between 1989 until 2009, and then declined steadily between 2009 and 2018. Average annual growth increments of huronicus lake charr were smallest in 2015-2018. For lean lake charr, average annual growth increments varied erratically from 1984 through 1990, declined from 1991 through 1995, increased from 1996 through 2009, and then declined between 2009 and 2018. Average annual growth increments of lean lake charr were nearly as small in 2015-2018 as in 1991-1995. Over the entire period, mean annual growth increments were 44% more variable for lean than for huronicus ecotypes (i.e., growth varied more among years for leans than huronicus overall; Table A2). From 2009 to 2018, mean increment width declined 20% faster for leans than huronicus (i.e., growth of leans declined faster after 2009 than growth of huronicus). Prior to 2009, mean increment width was 76% higher for leans than huronicus (i.e., leans grew faster before 2009 than huronicus).

**Table 2.** Tests of fixed effects for differences between lake charr morphs (huronicus or lean), sample years (2007 or 2018), and sexes (males or females) from a linear mixed-effects model of annual sagittal otolith growth increments as a function of a fixed age effect (age of increment formation), random year effects (year of increment formation), and random fish effects (fish-to-fish variation in growth) sampled in Rush Lake.

| Morph | Effect | Df<br>Numerator | Df<br>Denominator | F-Ratio | P-Value |
| --- | --- | --- | --- | --- | --- |
| Both | Age | 34 | 1,927 | 718.5 | $\leq 0.01$ |
| | Ecotype | 1 | 1,927 | 16.9 | $\leq 0.01$ |
|  | Sex | 1 | 1,927 | 0.4 | 0.5 |
|  | Year | 1 | 1,927 | 0.08 | 0.8 |
| Huronicus | Age | 30 | 1,182 | 515.4 | $\leq 0.01$ |
|  | Sex | 1 | 1,182 | 0.05 | 0.8 |
|  | Year | 1 | 1,182 | 0.9 | 0.3 |
| Lean | Age | 34 | 682 | 255.9 | $\leq 0.01$ |
|  | Sex | 1 | 682 | 1.4 | 0.2 |
|  | Year | 1 | 682 | 0.6 | 0.4 |

**Fig. 6.**
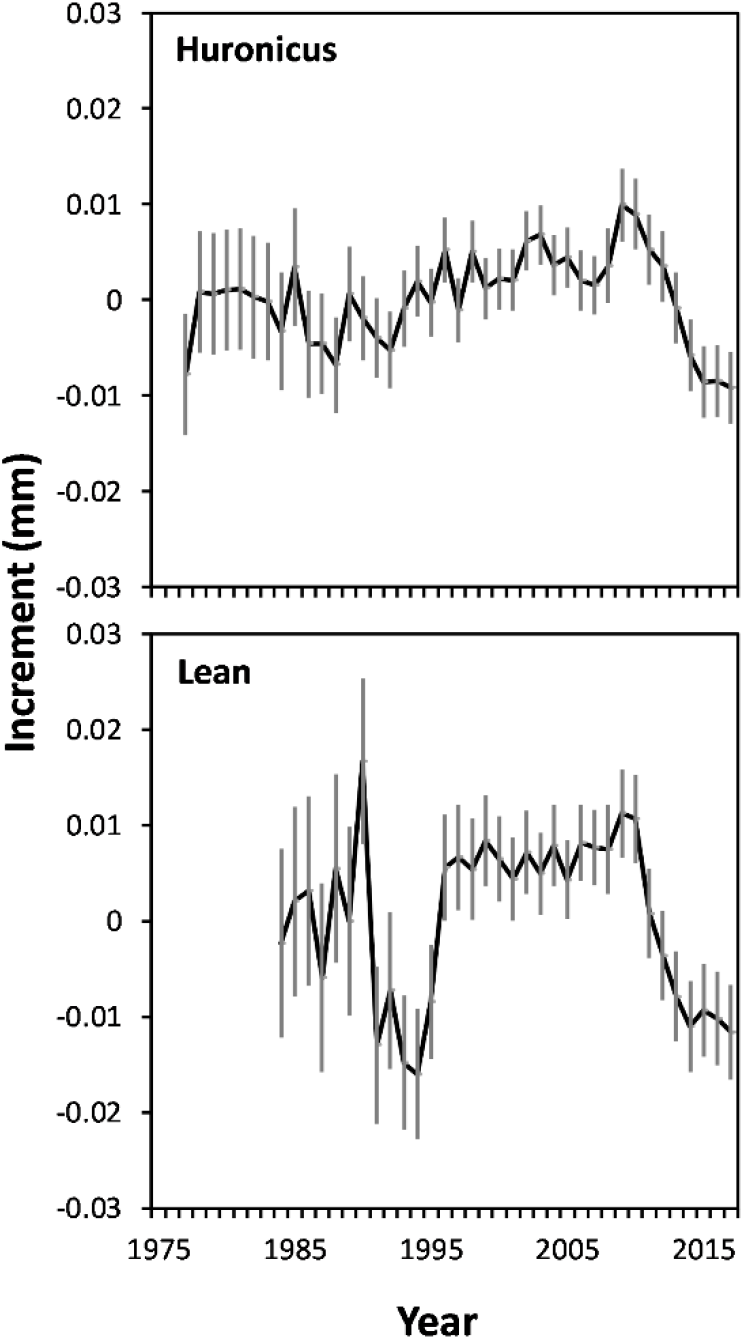
Annual growth increments (mm; random year effects from a linear mixed-effects model that also included fixed age effects and random fish effects; Weisberg et al. 2010) by calendar year for huronicus and lean lake charr ecotypes sampled from Rush Lake, in 2007 and 2018.

### Otolith increment cross-dating: growth of lake charr in relation to environmental variation

Inter-annual climate variation, particularly when including lagged effects, was correlated with fish growth for both ecotypes, as demonstrated by Pearson correlations regularly exceeding 0.2 (Fig. 7). For both lake charr ecotypes and year-corresponding and lagged effects, annual otolith growth increments were positively correlated with summer air temperatures (except for the lean corresponding year, slightly negative) and fall precipitation (i.e., more growth with warmer temperatures and more precipitation) and negatively correlated with summer and fall cloud cover (i.e., more growth with less cloud cover). In comparison, the direction of the relationship with winter to spring temperatures and precipitation as snow differed between the two ecotypes (Fig. 7 and A3). For each set of seasonal variables, inclusion of weighted climate data from previous years, to accommodate lagging effects, tended to strengthen correlations. Overall, based on the forward selection multiple regression models, the total amount of variation in otolith annual growth increments explained by climatic variables was greater for the lean ecotype than huronicus ecotype (R^2^ = 0.56 vs 0.35; Table A3). These regression models confirmed that growth increments of huronicus ecotypes were most strongly associated with winter to spring temperatures and precipitation as snow and secondarily with summer temperature, whereas the lean ecotype was most strongly associated with winter to spring temperatures and previous fall precipitation (Table A3). Among-ecotype differences in the relationship of annual growth increment with winter and spring temperatures and with precipitation as snow were then examined more closely, where regression analyses confirmed these differences (Fig. 7 and A3). Specifically, differences in annual growth increments between ecotypes for any given year indicated that cold and snowy winters tended to favor growth for the huronicus ecotype whereas warmer winters with less snow favored growth for the lean ecotype (Fig. 7 and A3).

**Fig. 7.**
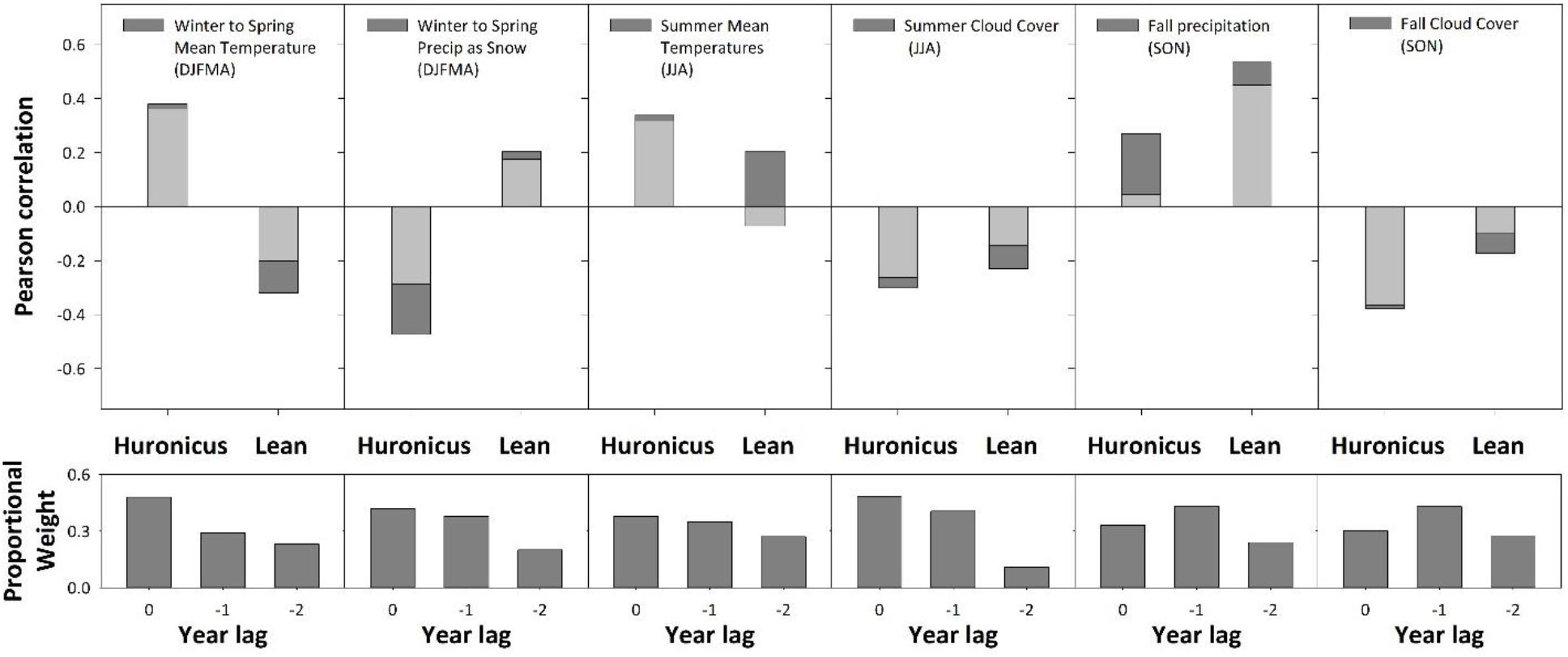
Pearson correlation coefficients between ARSTAN-detrended otolith growth increments and selected seasonal climate variables for each ecotype. Results include data combined across fish sampled in 2007 and 2018. Correlation values for each ecotype that used a proportional weighting scheme for seasonal variables across the current and two previous years is represented by dark bars, whereas light bars indicate correlation values with no weighting. Panels including proportional weight values (i.e., sum of weights equal to one) whereby the combination of weights was optimized to maximize correlation values shown in dark bars within the panel immediately above. Year lags correspond to the current year = 0, one previous year = −1 and two previous years = −2).

For the early life-stages of both ecotypes, defined here as ages 1-3, annual growth increments were correlated with cloud cover only. A negative relationship between early life stage growth and summer cloud cover (i.e., more growth with less cloud cover) occurred for both the huronicus (R^2^ = 0.19, p = 0.01) and the lean (R^2^ = 0.43, p < 0.01; Fig. 8) ecotypes. Annual growth of early life-stage lake charr appeared to be correlated with the temporal trend of summer cloud cover. For example, cloud cover in July has decreased by up to 33% over the past three decades, resulting in higher annual growth in early life-stage lake charr from 2007 to 2018 (Fig. 8).

**Figure 8.**
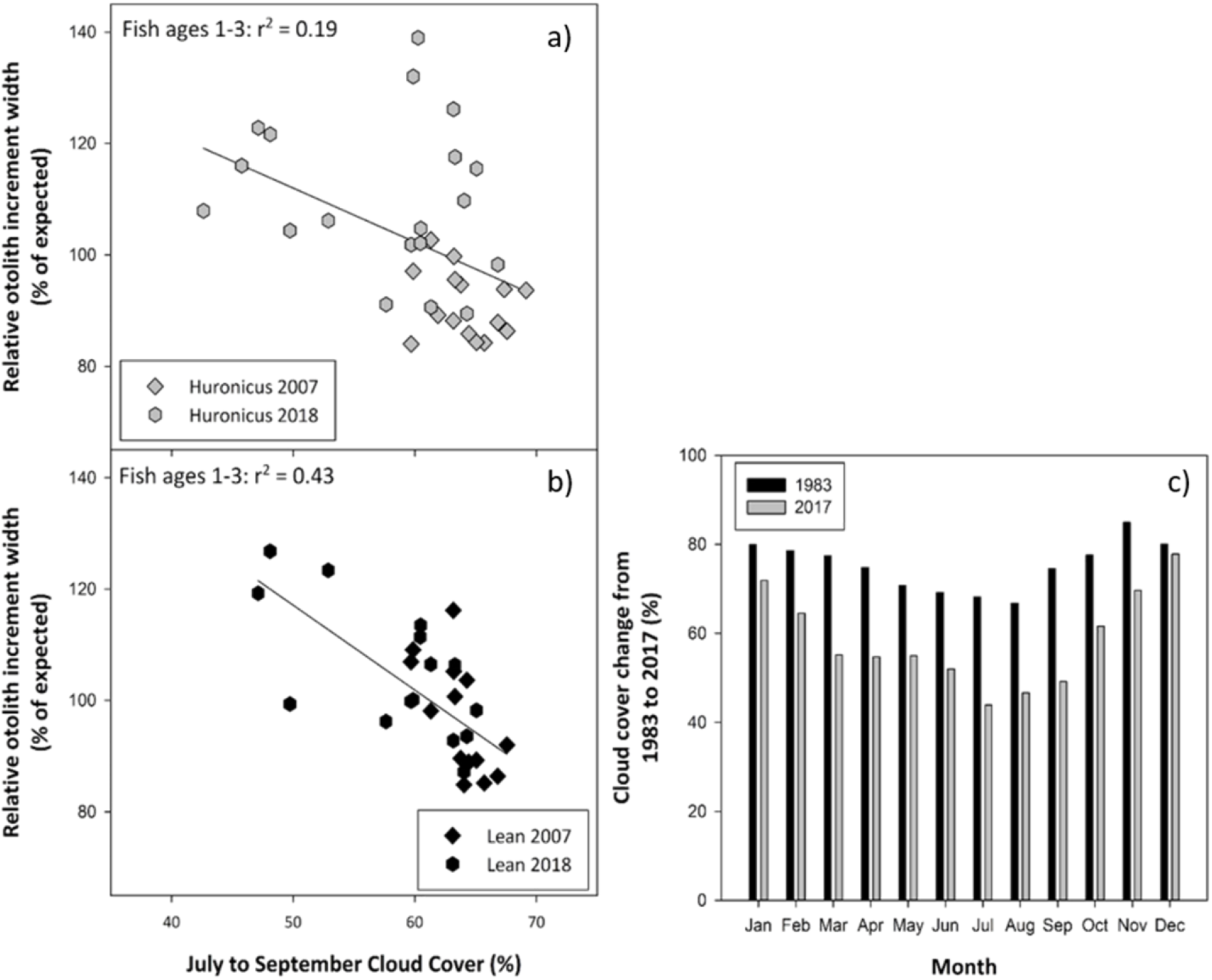
Otolith increment growth variation among ecotypes and collection periods for ages 1-3 plotted versus regional cloud cover from July to September for the same year without weighting or lag-effects included in a) and b). Data were detrended using regional chronology standardization (see Methods and Table A1). Within a given year, ecotype and collection period, otolith growth data for early stage growth were averaged across ages 1-3. In c), reductions in regional cloud cover by month from 1983 to 2017. Shown here are the predicted values for 1983 and 2017 using linear regressions fitted to interannual cloud cover data by month across the same time period. These two years represent endpoints for which otolith data were replicated enough to allow chronology construction. Cloud cover data were from airports within 7 km of Lake Superior since the lake itself strongly controls nearby weather (see Voelker et al. 2019).

## Discussion

In this study, we demonstrated a rapid phenotypic shift that occurred for two lake charr ecotypes in a small lake. Within the last decade, a major decline in annual growth increments has occurred with a corresponding morphological shift in body and head shape for both lean and huronicus ecotypes. Even though the lean (shallow-water) ecotype displayed a greater annual growth variation over the years than the huronicus (deep-water) ecotype, both ecotypes displayed similar magnitudes of morphological change in the most recent decade resulting in analogous “sensitivity” in phenotypic change, meaning “responsiveness” independent of habitats. We interpret these results to mean that the response threshold that determines individual sensitivity to a particular cue (i.e., environmental variables) caused similar phenotypic changes by individuals of both ecotypes (Moczek 2010; Baerwald *et al*. 2016), in spite of differences in sensitivity between growth responses of the two ecotypes. A rapid phenotypic shift was somewhat surprising because mobile predators, such as lake charr, they exploit a large range of prey, and thus, can compensate rapidly to prey fluctuations and dampen oscillations or fluctuations of lower trophic levels (McCann *et al*. 2005; McCann 2012) being caused by environmental variables.

One assumption often made in relation to intraspecific diversity, mostly tested experimentally or modelled, is that a stable or predictable environment interacts with underlying variation in expression of phenotypes (Wagner & Schwenk 2000; Svanbäck *et al*. 2009; Westneat *et al*. 2015; Skúlason *et al*. 2019). In our study, the magnitude and direction of the observed phenotypic shift in both annual growth and morphology over a single decade was consistent for each ecotype and suggested similar pathways to which phenotypic variation was expressed. The degree of phenotypic variation that occurs within an ecotype theoretically depends on the relative strength and time of mechanisms that drive phenotypic change (Wood *et al*. 2020), the environment is not necessary constant across cohorts (Johnson *et al*. 2014). In our case, the observed phenotypic shift was relatively small, but nonetheless, detectable (i.e., cryptic eco-evolutionary dynamics; Kinnison *et al*. 2015). Several questions arise from our results, but one of interest is the organism’s capacity for phenotype acclimation to a changing environment (via phenotypic plasticity and adaptation; Huey & Berrigan 1996; Gorsuch *et al*. 2010). Was this phenotypic shift an isolated event or does this type of change occur frequently in this system? Answers to the question of phenotypic acclimation and the frequency of its occurrence within and among systems would require long data sets collected over multiples decades and would help to fill an important knowledge gap about non-equilibrium population dynamics affecting evolutionary dynamics.

Biological complexity is a key component of ecological variability and evolution (McShea & Brandon 2010; McShea 2017). Complexity is a recurring idea in evolutionary biology that can refer to a number of concepts at different levels (organism to ecosystem), but biological complexity is defined herein as mechanisms producing (or limiting) variation in phenotypic expression (e.g., morphology; Heim *et al*. 2017; Duclos *et al*. 2019). In this study, a decline in annual growth increment for lean and huronicus ecotypes occurred at the same time as a consistent phenotypic shift in morphology and these changes occurred within the time frame of a decade. Growth rate in fishes has been shown to drive intraspecific morphological differentiation (Tonn *et al*. 1994; Olsson *et al*. 2007; Chivers *et al*. 2008). Growth rate seems to play a key role in regulating morphological expression, but underlying mechanisms remain uncertain (Olsson *et al*. 2006; Svanbäck *et al*. 2017; Franklin *et al*. 2018). A possible explanation is that at higher growth rates, energy is allocated to somatic growth and morphology modulation in addition to metabolic maintenance, but that at lower growth rates, energy is used almost exclusively for metabolic maintenance and/or reproduction (Olsson *et al*. 2006; Svanbäck *et al*. 2017). Our results concur with this latter mechanism because lake charr sampled in 2007 had higher annual growth rates and less morphological overlap between ecotypes than lake charr sampled in 2018. An organism’s response to change (abiotic and biotic) can include variation in the mean phenotype itself, but it can also include differences in the phenotypic variance (O’Dea *et al*. 2019). To our knowledge, few other field studies have demonstrated a temporal change in morphology and growth rate, within an ecotype much less, temporal changes that were consistent between ecotypes (but see Svanbäck *et al*. 2009 for an example of consistent phenotypic variation, in magnitude and direction between ecotypes (PC1: 2003 to 2004)).

Many examples of taxa exist where population abundance fluctuates (e.g., density) over time as a result of variation in resource levels, often influenced by environmental changes (Grant & Grant 1992; Persson & De Roos 2003; Svanbäck *et al*. 2009). In this study, some aspects of body and head shape shifts may have been mediated by changes in growth rates and body condition, which likely, were induced by variation in food availability and abiotic conditions. Body condition can affect body shape in fish, and often reflects bulkiness of individuals, and consequently body depth (Olsson *et al*. 2006; Borcherding & Magnhagen 2008; Jacobson *et al*. 2015; Svanbäck *et al*. 2017). Although both ecotypes were subjected to annual growth rate declines, the huronicus ecotype was more affected by body condition changes than the lean ecotype, which might suggest physiological differences between ecotypes in the dynamics of energy processing (e.g., metabolism and reproduction) and storage (Goetz *et al*. 2013). For example, in 2018, 30.8% of the huronicus females were in a resting reproductive stage compared to only 8.3% of lean females. In contrast to females, all males in 2018 of both ecotypes were reproductively mature (*Unpublished data*). Differences in energetic requirements associated to reproductive output appear related to the apparent skipped spawning patterns observed by ecotype and sex.

Spatial and temporal fluctuations of trophic resources in Rush Lake could have influenced how individuals used these resources and the resulting annual growth rate patterns (influenced by density-dependent fluctuations and intraspecific competition; Svanbäck *et al*. 2009; Jacobson *et al*. 2015). Such spatial and temporal periodicity can occur at different scales (e.g., for temporal periodicity: seasonal, inter-annual, decadal) and be driven by fluctuations in abiotic (e.g., temperature, precipitation, light, nutrients) and biotic processes (e.g., growth, reproduction, trophic interactions), thereby shaping species behavior and “rewiring” food webs (McMeans *et al*. 2015; Bartley *et al*. 2019). Developmental stability suppresses variation in phenotypic expression of a single genotype within a single environment, thus it can modulate the magnitude and structure of phenotypic variation among individuals (Willmore *et al*. 2006; Lazić *et al*. 2015). Organisms can be affected by a single perturbation of the timing or rate in development, which has been perceived to be a means to produce trait novelty (e.g., heterochrony; Parsons *et al*. 2011; Lazić *et al*. 2015; Westneat *et al*. 2015). Allometry, the shape variation associated with size variation (Zelditch *et al*. 2012), is also seen as a canalized process and an interacting agent that can limit morphological variation (Klingenberg 2010). At the very least, some of our results represent a case of plastic allometry for the lean ecotype in 2018. Considering that the lean body condition did not markedly differ between sampling years and were within the species range values (Hansen et al, In press), we are confident that morphological differences were not due to starvation of individuals. Altogether, the expression of phenotypic variation observed in this study was probably influenced by more than one varying niche dimension (e.g., temperature, resources level, competition) with concomitant effects (e.g., developmental stability and allometry).

Lake charr seem to show a fluctuating nature of phenotypic expression (adaptive or not), closer to Eurasian perch (*Perca fluviatilis*) than other salmonids such as Arctic charr (*Salvelinus alpinus*). Both lake charr and perch appear to show less discreteness between ecotypes than Arctic charr but yet, a higher level of phenotypic variance (Chavarie et al., accepted; Jonsson & Jonsson 2001; Svanbäck *et al*. 2009). When compared to lake charr ecotypes from other lakes, the genetic diversity and divergence of lake charr in Rush Lake is low for both ecotypes (Chavarie *et al*. 2016). This low genetic diversity and divergence favor the hypothesis that phenotypic variation is the result of phenotypic plasticity rather than genetic adaptations (although rapid genetic change cannot be excluded). Epigenetically mediated biological complexity is known to be an important process to tailor phenotypic reaction norms (e.g., linear and nonlinear) to selective environmental pressures (Crozier & Hutchings 2014; Ramler *et al*. 2014; Duclos *et al*. 2019). A rapid change in the environment can also induce changes in the phenotypic variance within an ecotype by exposing previously hidden cryptic genetic variation or by inducing new epigenetic changes (O’Dea *et al*. 2016). It has been hypothesized that heritable epigenetic mechanisms can lead to phenotypic variation generated by bet hedging strategies, whereas phenotypic variability buffers varying environments (O’Dea *et al*. 2016). Environmental factors can induce a component of variation that introduces a fined-grained variation around any coarse-scale temporal trends, resulting in year-to-year variation in phenotypes but not in genotypes – because genetic changes are not expected to be so fine-grained (Merilä & Hendry 2014). Thus, phenotypic variability, both in means and variance, can have evolutionary scope for a population in the face of changing selection regimes by affecting population dynamics and probabilities of extinction (Chevin *et al*. 2010; Reed *et al*. 2010; Johnson *et al*. 2014).

Synchronous change in growth rates of fishes have been observed at global scales, with major declines in growth linked to climate change (Thresher *et al*. 2007; Sheridan & Bickford 2011; Baudron *et al*. 2014). However, little is known of how growth trajectories and their associated phenotypic reaction norms that integrate environmental factors may differ within (e.g., cohorts) and among ecotypes in freshwater ecosystems (Heino *et al*. 2002; Johnson *et al*. 2014). In Rush Lake, similarities and differences of the annual growth rates of lake charr ecotypes were correlated to environmental variation. In our study, cloud cover was the main environmental variable that had steadily decreased over the same time period that lake charr growth declined (except for early stage) and morphology shifted in Rush Lake (Fig. 8). Environmental heterogeneity is thought to have stronger effects on morphology at early life stages (Johnson *et al*. 2014; Morris 2014; Ramler *et al*. 2014), suggesting that the effects of the cloud cover on lake charr might have been more significant at age 1-3 years than during later stages of life. The effect of cloud cover on growth at age 1-3 was stronger in the lean than the huronicus ecotype, which might explain why allometry was detected only for the lean ecotype in 2018. The relationship between lake charr annual otolith growth and cloud cover could be related to how solar irradiance can co-vary with other climate variables that may affect fish growth (Reist *et al*. 2006; Poesch *et al*. 2016). For example, in Lake Superior, cloud cover and light penetration were shown to affect the degree of heterotrophy (primary production < community respiration; Brothers & Sibley 2018). Rush Lake is located only two km away from Lake Superior at its nearest point and this proximity produces strong lake-effect climate phenomena during summer (Voelker *et al*. 2019). Primary production can have cascading effects to top-predators, thereby influencing the level and quality of available food ration (Hunter & Price 1992). Summer temperature and fall precipitation were also positively associated with growth rates. When higher temperatures are accompanied by suitable net addition of food ration (e.g., from direct and indirect effects of temperature and precipitation factors), increases in growth could be expected up to the point of the optimum temperature of the species (Elliott & Hurley 2003; Elliott & Elliott 2010).

Response to winter climate appeared to vary between ecotypes, with environmental variables (e.g., temperature and precipitation as snow), correlated in different directions with growth rates (directly or indirectly). Why the deep-water ecotype (huronicus) would show higher annual growth increments in years with warm winter temperatures and low snow cover (via direct or lagging effects) and the shallow-water ecotype (lean) would have higher annual growth increments in years with cold winter temperature and high snow level is unknown. These results could be explained in part by reduced habitat partitioning between ecotypes during winter, along with an increase of intraspecific competition, affecting energy storage (Amundsen *et al*. 2008). Another explanation, which is not exclusive of the previous one, could be that each ecotype’s prey types, density and quality (e.g., time response to environmental variable) is modified differently by lagging effects from winter environmental conditions (e.g., ice cover duration, ice and snow thickness). The lean ecotype is known to feed on forage fish whereas the huronicus ecotype mainly feeds on the invertebrate *Mysis* (Chavarie et al. 2016b). Invertebrates can differ in response time and magnitude to environmental changes compared to forage fishes (Heino et al. 2009, Wrona et al. 2016, Wrona et al. 2006). This hypothesis of a differential response of prey items to environmental conditions seems to be supported by correlations strengthened with lake charr growth when weighted climate data from previous years were included. This result might be expected for an organism in which growth rates may draw on a mixture of recently acquired and stored resources or where climate variables in one year may affect the abundance and composition of prey in subsequent years. For the most part, winter environmental conditions can play an essential role in ecological and evolutionary processes that define life-history characteristics (e.g., somatic growth, size and age-at-maturity, reproduction investment, and longevity) of lacustrine species (Shuter *et al*. 2012).

## Conclusion

We demonstrated such rapid morphological change in Rush Lake lake charr, a change that was expressed similarly by both lean and huronicus ecotypes over a decade of time. These shifts in morphology shared by both ecotypes were synchronous with declining in cloud cover over the same time frame. The mechanisms that connect annual growth increments with morphological modulation are not fully understood (Olsson *et al*. 2006; Olsson *et al*. 2007; Svanbäck *et al*. 2009); however, the biology underlying phenotypic variation can have major implications for populations responding to climate change. Multidimensional phenotypic variability and its influence on population dynamics patterns is a relatively poorly studied phenomena (Westneat *et al*. 2015), but individual and population resistance and resilience to climatic changes may depend on this variability (Johnson *et al*. 2014). Mean fitness can improve more rapidly through plasticity than through genetic adaptation (at least on the short-term perspective; Hendry *et al*. 2011).

The similarity in phenotypic response expressed by both ecotypes raises the question whether organisms in small lakes are more vulnerable to climate change than those in large lakes. Small lakes generally sustain a higher degree of habitat coupling (e.g., littoral-pelagic; Schindler & Scheuerell 2002; Dolson *et al*. 2009), which is critical to food-web dynamics. Thus, the degree of habitat coupling found in each freshwater ecosystem might translate to its degree of vulnerability to climate change. Field studies, such as ours, that focus on temporal phenotypic instability within an aquatic ecosystem promise to clarify our understanding of how the interplay among phenotypes, trophic dynamics, and environmental context influences both ecosystem and evolutionary processes (Ware *et al*. 2019).

## Acknowledgements

We thank the Huron Mountain Club for access to their lands, housing, and lakes and for sharing their knowledge of the lake charr of Rush Lake. Special thanks to Kerry Woods, Director of Research, Huron Mountain Club Wildlife Foundation for coordinating and supporting the project. Financial support was provided by the Great Lakes Fishery Commission. Any use of trade, firm or product names is for descriptive purposes only and does not imply endorsement by the U.S. Government. The findings and conclusions in this article are those of the authors and do not necessarily represent the views of the US Fish and Wildlife Service.

## Appendix

**Table A1.** Listing and comparison of analytical methods used to analyze otolith growth data for different purposes and associated results.

| Analytical method for otolith growth | Advantages | Disadvantages | Results |
| --- | --- | --- | --- |
| Non-linear mixed-effects modeling | Uses all data, so it can be used with all fish morphology measurements to back-calculate fish growth and asymptotic length | Cross-dating not confirmed, more environmental noise not related to climate drivers | Table 1, Table 2, Fig. 5, Fig. 6, Table A1 |
| Cross-dated and ARSTAN-detrended | Conventional approach for identifying climate drivers of inter-annual variation in long-lived organisms such as trees | Short age span of fish excludes identification of decadal or greater environmental drivers (i.e., "segment length curse"). Some loss of data when otolith time series do not cross-date | Fig. 7, Table A2, Fig A3. |
| Cross-dated and RCS-detrended | Alternative approach for identifying climate drivers that preserves decadal and longer signals | Not as efficient at accurately extracting inter-annual variation compared to ARSTAN-detrending | Fig. 8, Fig A3 |

**Table A2.** Difference in annual growth increments (mm; random year effects from a linear mixed-effects model that also included fixed age effects and random fish effects; Weisberg et al. 2010) by calendar year for huronicus and lean lake charr ecotypes sampled from Rush Lake, in 2007 and 2018 (Figure 6).

| Metric | Huronicus | Lean | % |
| --- | --- | --- | --- |
| SD | 0.0049 | 0.0088 | 44 |
| Slope | -0.002 | -0.0026 | 20 |
| Decline | 0.018 | 0.024 | 24 |
| Mean | 0.0004 | 0.0018 | 76 |

**Table A3.** Forward selection multiple regression modeling of detrended otolith increment growth chronologies for lake charr ecotypes (huronicus or lean) sampled in Rush Lake in 2007 and 2018, as predicted by weighted seasonal climate variables. Environmental variables are represented as follow: Twin = winter temperature, SNOwin = winter snow, Tsum = summer temperature, and Pfall = fall precipitation.

|  | **Models** | **R<sup>2</sup>** | **P-value** | **AIC** |
| --- | --- | --- | --- | --- |
| **Huronicus** | Twin | 0.23 | 0.01 | -90.3 |
|  | SNwin | 0.26 | <0.01 | -91.5 |
|  | SNwin + Tsum | 0.35 | <0.01 | -93.1 |
| **Lean** | Twin | 0.31 | <0.01 | -59.6 |
|  | Pfal | 0.37 | <0.01 | -61.5 |
|  | Twin + Pfal | 0.56 | <0.01 | -67.7 |

**Fig. A1.**
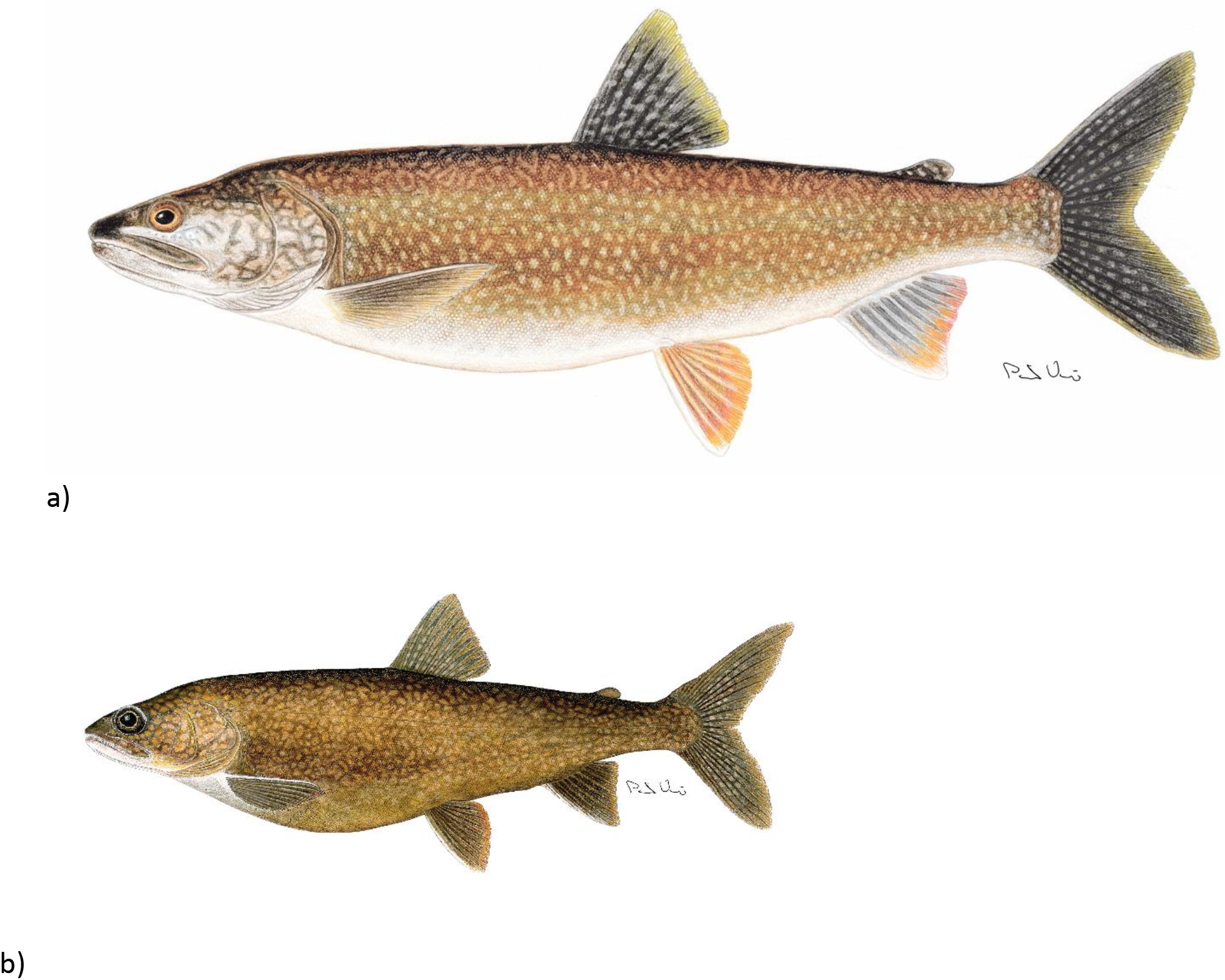
Lean-like (a) and huronicus (b) lake charr *Salvelinus namaycush* ecotypes of Rush Lake. Illustration by P.Vecsei, after Chavarie *et al*. (2016)

**Fig. A2.**
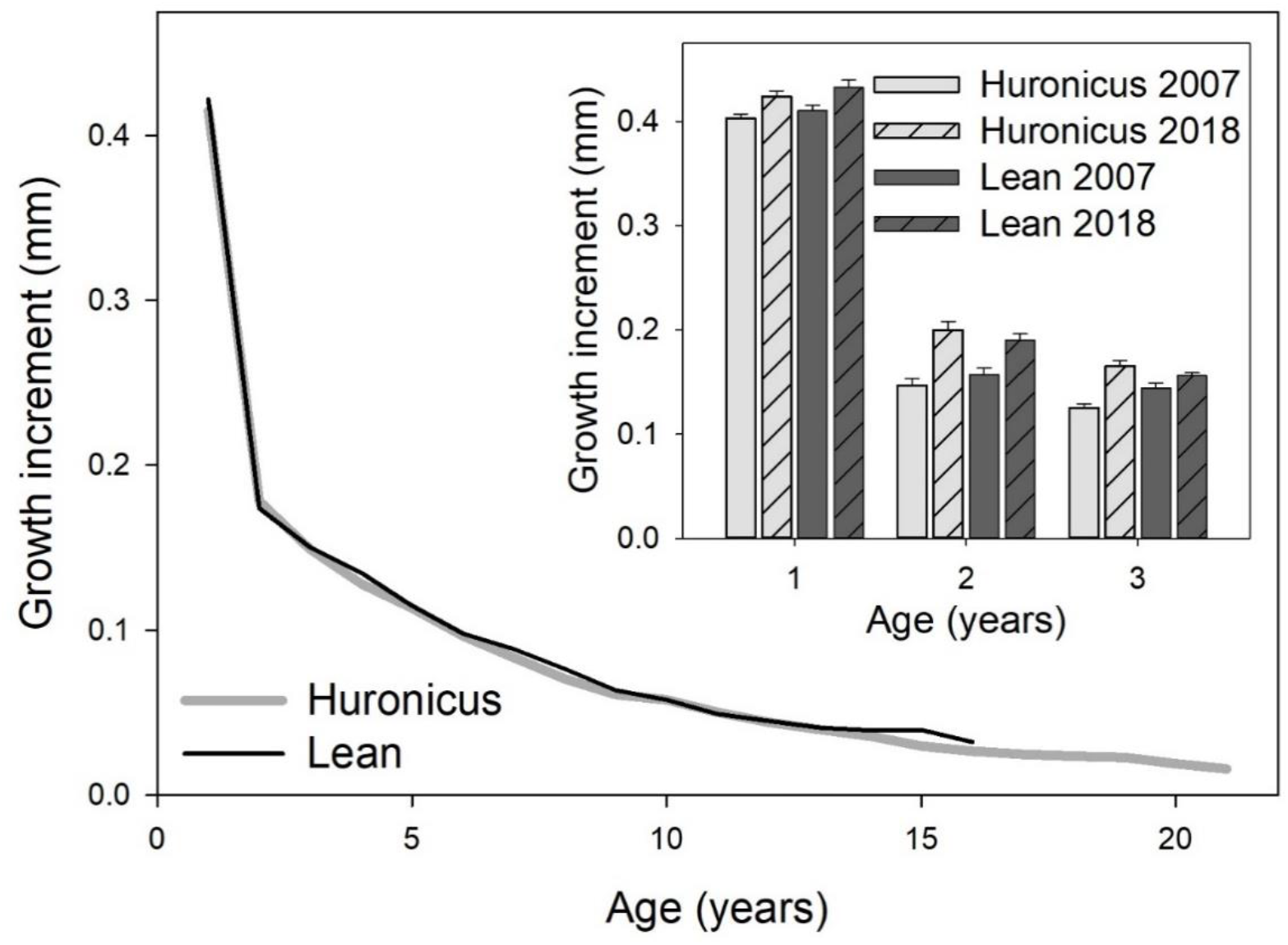
Otolith growth increment plotted by age between lake charr ecotypes (lines) and detailed across the first three years for lake charr morphs (huronicus or lean) sampled in 2007 and 2018.

**Fig. A3.**
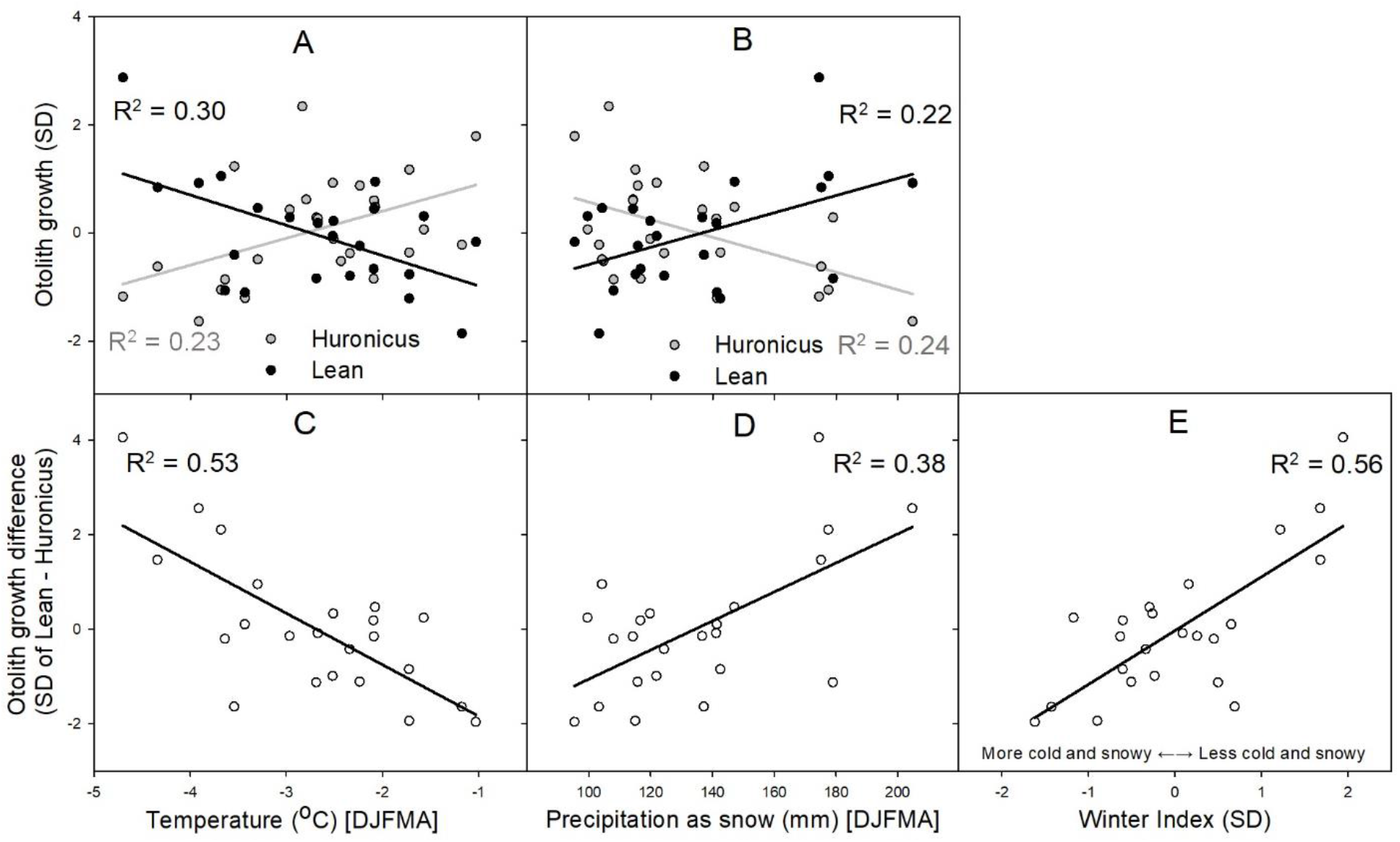
Standard deviations (SD) of ARSTAN-detrended growth increment versus previous winter temperatures and precipitation as snow. Growth SD between lake charr ecotypes sampled in Rush Lake in 2007 and 2018, plotted versus weighted winter climate variables (A, B), SD of the difference between ecotype growth versus weighted winter variables (C, D) and SD of the difference between ecotype growth plotted versus a “winter index” that averaged the SD of sign-reversed temperature and precipitation as snow.

